# The role of clathrin in exocytosis and the mutual regulation of endo- and exocytosis in plant cells

**DOI:** 10.1101/2021.11.17.468992

**Authors:** Maciek Adamowski, Ivana Matijević, Jiří Friml

## Abstract

Within the plant endomembrane system, the vesicle coat protein clathrin localizes to the plasma membrane (PM) and the *trans*-Golgi Network/Early Endosome (TGN/EE). While the role of clathrin as a major component of endocytosis at the PM is well established, its function at TGN/EE, possibly in exocytosis or the vacuolar pathway, is a matter of debate. This shared function of clathrin also opens a question whether plant cells possess a homeostatic mechanisms that balance rates of opposite trafficking routes, such as endo- and exocytosis. Here we address these questions using lines inducibly silencing *CLATHRIN HEAVY CHAIN* (*CHC*). We find a relocation of exocytic soluble and integral membrane protein cargoes to the vacuole, supporting a function of clathrin in exocytosis. A comparison with lines overexpressing AUXILIN-LIKE1, where inhibition of CME precedes rerouting of secretory cargoes, does not support a homeostatic regulatory mechanism adjusting exocytosis to the rates of endocytosis. Complementary experiments reveal only minor and variably detectable reductions in the rates of CME in secretory mutants, also not indicative of a converse homeostatic mechanism adjusting rates of endocytosis to the rates of secretion.

## Introduction

The endomembrane system of plant cells serves multitudes of cellular functions; not least, it is responsible for the biosynthesis, processing, and distribution of cell surface proteins, such as receptors, channels, and transporters, and of most carbohydrate cell wall components. The endomembrane system is organized into distinct compartments, where particular biochemical processes occur, and sorting decisions take place. The endomembrane compartments are linked by specific trafficking processes, which carry cargoes from one compartment to another. Trafficking through the endomembrane system often involves formation of small carriers - cargo-loaded vesicles - which bud off one compartment and fuse with another (Buchanan et al., 2015). In other instances, trafficking occurs through a gradual maturation of one compartment type into another (Scheuring et al., 2011; Day et al., 2013).

One key element of the endomembrane system is the secretory pathway, executing the fundamental function of the delivery of proteins and cell wall components to the cell periphery. This pathway originates at the endoplasmic reticulum (ER), and passes typically through the Golgi apparatus (GA) and the trans-Golgi Network (TGN) to finally reach the plasma membrane (PM) and the apoplast by a final step of vesicular transport termed exocytosis. Alternative secretory pathways have been described as well (Wang et al., 2017; Žárský et al., 2009). In turn, retrograde traffic from the PM, termed endocytic traffic, internalizes PM-localised and apoplastic components which may be then recycled through exocytosis, or directed, through early and late endosomes, towards a terminal offshoot of the endomembrane system in the vacuole, where protein degradation occurs. The initial step of the endocytic pathway is a vesicular transport process termed endocytosis. The major form of endocytosis in plants relies on the coat protein clathrin (Dhonukshe et al., 2007), and as such, is named clathrin-mediated endocytosis (CME; reviewed in Reynolds et al., 2018). A multitude of other factors participates in CME and their characterization is an active field of study.

Beside the PM, clathrin is present at the trans-Golgi Network/Early Endosome (TGN/EE) compartment (Ito et al., 2012; Kang et al., 2011). The function of clathrin at this compartment is usually assigned to the formation of clathrin-coated vesicles (CCVs) with cargoes destined for the vacuolar route (Buchanan et al., 2015; discussed in Robinson and Pimpl, 2014). Exocytosis, in turn, is thought to occur through secretory vesicles (SV) formed without protein coats (Žárský et al., 2009). Yet, indirect lines of evidence link clathrin to the exocytic process. During clathrin-coated pit (CCP) formation, the clathrin coat is physically linked with the membrane of the nascent vesicle by adaptors (Owen et al., 2004). At the TGN compartment, CCVs are formed with the aid of Adaptor Protein 1 (AP-1) complex (Wang et al., 2013; Park et al., 2013; Robinson and Pimpl, 2014). The AP-1 complex is, in turn, recruited to membranes as an effector of activated ARF small GTPases, as indicated by studies on non-plant homologues (Paczkowski et al., 2015). In *A. thaliana*, loss-of-function mutants of *AP1M2*, encoding an AP-1 subunit, exhibit deficiencies in secretion of soluble proteins (Park et al., 2013). Inhibition of TGN-localized ARF activators, the BIG family of ARF-GEFs (Guanine nucleotide Exchange Factors for ARFs), similarly leads to an arrest of secretion, concomitant with a relocation of AP-1 into the cytosol (Richter et al., 2014). Together, these results are consistent with a scenario where AP-1-containing CCVs formed with the aid of ARFs activated by BIG ARF-GEFs, participate in exocytosis from the TGN. Yet, *ap1m2* mutants exhibit defects of vacuolar traffic as well, and it may be that either the secretory or the vacuolar trafficking defects in *ap1m2* are indirect consequences of a malfunctioning TGN compartment, rather than representing genuine pathways mediated by vesicles coated with AP-1 and clathrin. In strong support of a function of AP-1 in vacuolar sorting, an AP-1-binding motif mediating vacuolar targeting has been described (Wang et al., 2014). Furthermore, while the deactivation of BIG ARF-GEFs does abolish the recruitment of AP-1 to the TGN (Richter et al., 2014), it is almost certain that TGN-localized ARFs recruit other effectors as well, and as such, the inhibition of exocytosis by BIG deactivation is not specifically informative about an exocytic role of AP-1 and by extension, of clathrin. Additionally, evidence from loss-of-function mutants of an *A. thaliana* AP-3 adaptor complex, which in immunoprecipitation experiments binds clathrin, are too, indirectly indicative of a function of clathrin in the vacuolar pathway (Zwiewka et al., 2011; Feraru et al., 2010). In turn, loss of function mutants of TGN-localized monomeric clathrin adaptors MODIFIED TRANSPORT TO THE VACUOLE1 (MTV1) and EPSIN1 indicate clathrin functions in both vacuolar, and secretory, pathways (Sauer et al., 2013; Heinze et al., 2020; Collins et al., 2020). Finally, multivesicular bodies (MVBs), organelles acting as late endosomes in delivery of cargo to the vacuole, were observed to directly mature from the TGN compartment, and soluble vacuolar cargo to reach the vacuole following inhibition of clathrin function, arguing for a lack of necessity of clathrin-mediated vesicular transport for vacuolar delivery (Scheuring et al., 2011). Taken together, the accumulated data warrants a closer investigation of the function of TGN-localized clathrin.

Another, potentially linked question is that of a mutual relationship of trafficking processes within the endomembrane system. The endomembrane system mode of operation, which includes transport in membrane-bound vesicles produced from one compartment and fusing with another compartment, leads to the movement of lipid membrane components as a result of cargo trafficking activity. The major site of membrane lipid biosynthesis is the ER (van Meer et al., 2008), and as such, this compartment can be seen as the source of both cargo and membrane within the endomembrane system. At the opposite end, the PM should contain, through the difference between membrane addition in secretory vesicles and membrane removal in endocytic vesicles, just the right amount of membrane to accommodate for cell growth, while retaining optimal membrane tension. The intermediate intracellular compartments, too, presumably need to be maintained in a balance between membrane reception with incoming vesicles and its removal due to vesicle production, or maturation. Therefore, it is conceivable that the rates at which distinct vesicular trafficking processes within a cell operate might be mutually dependent. The question of so understood “trafficking homeostasis” has been a matter of some research, especially focused on a potential regulation acting between endocytosis and exocytosis. Following the isolation of yeast *sec* mutants, deficient in secretory traffic (Novick et al., 1980; Spang, 2015), it was investigated whether the mutants have functional endocytosis. Some of the *sec* mutants, in particular those deficient in late steps of exocytosis, were indeed defective in an endocytic uptake of a fluorescent marker dye (Riezman, 1985). The yeast Rab GTPase Sec4p has been more recently described as a secretory component triggering endocytic events following delivery to the PM, and as such coupling the two processes (Johansen et al., 2016). The question of PM turnover in plant cells as a result of secretion and endocytosis was reviewed as early as 1988 (Steer, 1988), while the processes that may require coupling of endo- and exocytosis, and potential molecular players involved, were discussed in a recent review (Zhang et al., 2019). Interestingly, Larson et al. (2017) report that *clathrin heavy chain1* (*chc1*) mutants, deficient in a component of the clathrin coat, are defective in secretion of transiently expressed secretory yellow fluorescent protein (secYFP) while *syntaxin of plants 121* (*syp121*), a mutant deficient in a Qa-SNARE (for SNAP RECEPTOR) required for exocytosis, exhibits reduced internalization of an endocytic marker dye FM4-64. A recent study further supports a bidirectional mutual regulation between post-Golgi trafficking, and CME (Yan et al., 2021). Shifts in the relative rates of endocytosis and exocytosis in response to changes in osmotic conditions were described as well (Zwiewka et al., 2015). These results are indicative of mechanisms for mutual regulation between endocytosis and exocytosis.

In the present work, we further explore these two linked questions: that of a potential mutual regulatory relationship between endocytosis and exocytosis, and that of a potential function of clathrin in exocytosis. To this end, we make use of novel transgenic lines expressing inducible artificial microRNA (amiRNA) targeting *CLATHRIN HEAVY CHAIN* (*CHC*), in which we investigate effects on secretion of soluble and integral membrane protein cargoes. We further compare silencing of clathrin with effects of inducible overexpression of a putative uncoating factor for endocytic CCVs, AUXILIN-LIKE1, previously described as leading to inhibition of CME (Adamowski et al., 2018). Finally, to address the impact of the secretory pathway on endocytosis, we analyze CME in several mutants with deficient secretion with the use of Total Internal Reflection Fluorescence (TIRF) microscopy on fluorescent reporters for endocytic CCPs.

## Results

### Generation of inducible lines silencing *CHC*

To address directly the function of clathrin in secretion, we generated amiRNA lines targeting genes encoding CHC, which together with CLATHRIN LIGHT CHAIN (CLC) comprise the vesicular clathrin coat. We cloned three different amiRNA sequences, named *amiCHCa, amiCHCb*, and *amiCHCc*, all of which target both *CHC* homologues in *A. thaliana, CHC1* and *CHC2*. We expressed the amiRNAs in stably transformed *A. thaliana* plants under the control of the β-estradiol-inducible expression system. *XVE»amiCHCa, XVE»amiCHCb*, and *XVE»amiCHCc* constructs were introduced into the *secRFP* marker line (Samalova et al., 2006), which expresses a variant of red fluorescent protein (RFP) with a signal peptide, leading to its insertion into the ER and subsequently, secretion to the apoplast via the secretory pathway.

When cultured *in vitro* on media supplemented with β-estradiol, *XVE»amiCHCa*, *XVE»amiCHCb*, and *XVE»amiCHCc* seedlings exhibited greatly diminished rates of growth and development. After 6 d of growth, the seedlings were of minute size, usually without appreciable outgrowth of the root, with small cotyledons without green pigmentation (Figure 1A, B and Suppl. Figure 1A). When seedlings of these lines were germinated on standard media, transferred onto media supplemented with β-estradiol at day 3, and their root apical meristems (RAMs) observed under a light microscope 48 h later, the cells in the RAM were found to be enlarged, and characterized by large, swollen vacuoles (Figure 1C and Suppl. Figure 1B). This morphological characteristic was similar to that observed previously in lines inducibly overexpressing AUXILIN-LIKE1/2 (Adamowski et al., 2018). Downregulation of *CHC1* and *CHC2* expression after 48 h of transgene was confirmed by qPCR in lines expressing each amiRNA variant (Suppl. Figure 2).

**Figure 1.**
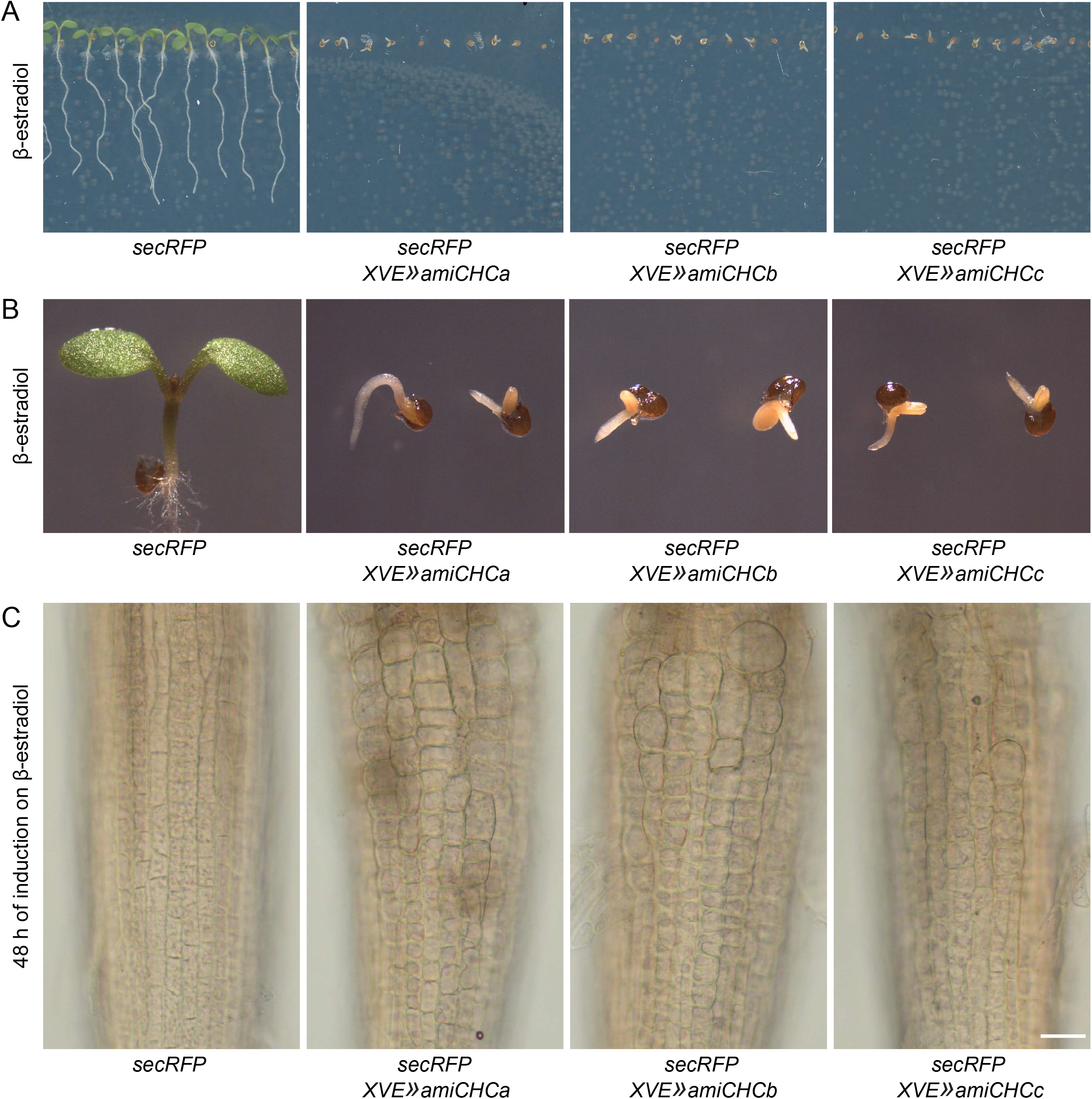
Morphology of *XVE»amiCHC* seedlings. (A) *secRFP XVE»amiCHC* lines grown on a medium supplemented with β-estradiol for 6 d. Silencing of *CHC* leads to a severe delay in development. Mock controls in Suppl. Figure 1A. (B) Detail of seedlings shown in (A). (C) Microscopic images of RAMs of *secRFP XVE»amiCHC* lines after 48 h of induction by β-estradiol. The RAMs of *XVE»amiCHC* lines exhibit enlarged cells with swollen vacuoles. Mock controls in Suppl. Figure 1B. Scale bar – 10 μm.

In summary, interference with clathrin function by inducible silencing of *CHC* lead to a severe inhibition of growth and development. In the following, we first characterize the consequences of *CHC* silencing on CME and on internal endomembrane compartments, and then, the consequences on secretion. For this characterization, we introduced the amiRNA-expressing constructs into several other fluorescent marker lines. Considering the overall frequency and severity of induced phenotypes among all raised *secRFP XVE»amiCHC* lines, we selected *XVE»amiCHCa* for further study, as expressing the most effective amiRNA sequence.

### Disruption of CME by silencing of *CHC*

To assess the effect of silencing of *CHC* on CME, and as such, additionally verify the correct function of the amiRNA transgene, we introduced *XVE»amiCHCa* into marker lines expressing fluorescent protein fusions of several main components of CME. We analyzed CLATHRIN LIGHT CHAIN2 (CLC2; Konopka et al., 2008), an isoform of CLC; TPLATE, a subunit of the TPLATE COMPLEX, functioning in plant CME presumably as an adaptor protein complex (Gadeyne et al., 2014); AP2A1, a subunit of the AP-2 adaptor complex (Di Rubbo et al., 2013); and DYNAMIN RELATED PROTEIN 1C (DRP1C; Konopka et al., 2008), a plant-specific isoform of dynamin, a mechanoenzyme acting in the separation of nascent CCV from the PM. In this and following sections, *amiCHC* constructs were induced for approximately 48 h prior to fluorescent live imaging.

When assessed by Confocal Laser Scanning Microscopy (CLSM) in seedling RAM epidermis, silencing of *CHC* lead to a relocation of CLC2-GFP from its variably detectable PM localization, into the cytosol (Figure 2A). The intracellular portion of CLC2-GFP also appeared to be lost from the TGN compartments (Figure 2A). Scarce remaining fluorescent spots were typically still observed, but these were likely autofluorescent structures, whose detection could not be avoided due to low expression of CLC2-GFP, as these signals could be still observed after bleaching of GFP by the application of high laser (Figure 2A, right). As will be seen below, TGN compartments, where clathrin localizes, are abundant in *XVE»amiCHCa*, as such presenting a different image. When assessed by TIRF microscopy in early elongation zones of the root (Figure 2B and Suppl. Movies 1 and 2), a strong, diffuse background signal of CLC2-GFP could be observed following *CHC* silencing, consistent with the observation by CLSM, but often, signals from scarce PM-localized CCPs, as well as TGN/EE compartments in the cytosol, were seen as well. In summary, down-regulation of *CHC* caused CLC, now unable to form triskelia with CHC, to partially relocate to the cytosol from its sites of action, both at PM and TGN.

**Figure 2.**
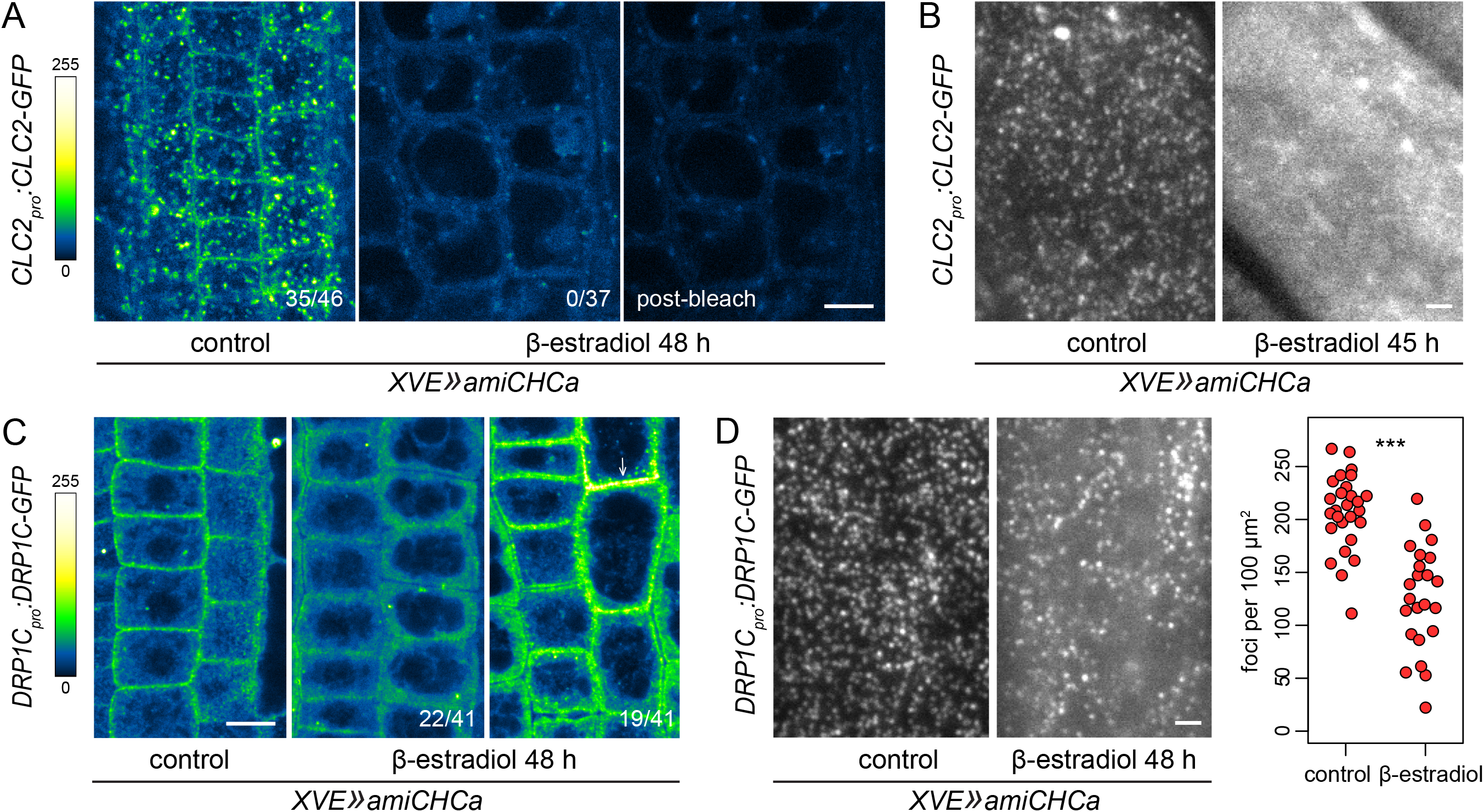
Effects of *CHC* silencing in *XVE»amiCHCa* on CLC2-GFP and DRP1C-GFP. (A) CLSM images of CLC2-GFP in RAMs of control and induced seedlings of *XVE»amiCHCa*. Down-regulation of *CHC* lead to relocation of CLC2-GFP to the cytosol. The sparse remaining signals appear to be autofluorescent structures, as they remain fluorescent after photobleaching of GFP (right). Numbers in bottom rights corner indicate the ratio of roots with PM signals. Scale bar – 10 μm. (B) TIRF images of CLC2-GFP in roots of control and induced seedlings of *XVE»amiCHCa*. Following silencing of *CHC*, CLC2-GFP is observed as a bright diffuse signal in the cytosol, with few CCPs and TGN-localized CLC2-GFP signals visible in some of the samples. Scale bar – 2 μm. (C) CLSM images of DRP1C-GFP in RAMs of control and induced seedlings of *XVE»amiCHCa*. Down-regulation of *CHC* lead to variable outcomes, with 22/41 seedlings exhibiting loss of PM localization of DRP1C-GFP (middle panel), and 19/41 retaining PM signals of normal or abnormally high intensity (right panel, arrow). Scale bar – 10 μm. (D) TIRF images of DRP1C-GFP in roots of control and induced seedlings of *XVE»amiCHCa*. Down-regulation of *CHC* caused a reduction in average density of DRP1C-positive foci at the PM. Graph shows collated quantifications of average foci density from all experiments, each data point represents a single root. Mock: 207 ± 35 foci per 100 μm^2^, n=27; β-estradiol: 125 ± 48 foci per 100 μm^2^, n=23. Values were compared using a *t* test, *** P≤0.001. Scale bar – 2 μm.

The effects of *CHC* silencing on DRP1C-GFP observed with CLSM in RAM epidermis were variable. The PM signal of DRP1C-GFP was decreased or absent in approximately half of the seedlings (Figure 2C, middle) while in the other half of the analysed population, the signals either remained normal, or, were clearly increased compared with the control (Figure 2C, right, arrow). TIRF microscopy in early elongation zones of the root indicated an approximately two-fold decrease in the average density of DRP1C-positive foci at the PM following down-regulation of *CHC* (Figure 2D), while cells with abnormally high densities of DRP1C-positive foci were not observed.

In turn, after silencing of *CHC*, the intensity of both TPLATE-GFP and AP2A1-TagRFP marker signal was consistently increased at the PMs of RAM epidermis, as captured using CLSM (Figure 3A and 3B). A similar observation was made in *XVE»AUXILIN-LIKE1* lines (Adamowski et al., 2018). It appears that similarly to this former case, interference with clathrin function by *amiCHCa* causes the early-arriving adaptor complexes to be strongly recruited into futile attempts of CCP formation. Yet, TIRF microscopy in early elongation zones of seedling roots did not reveal any increase in the average density of TPLATE-GFP-positive foci at the PM (Figure 3C). AP2A1-TagRFP was not analysed by TIRF microscopy due to low detection. The observations can be interpreted two-fold: The effect observed with CLSM may be limited to the meristematic zone, or, the effect does not represent an increase in the density of initiated CCP formation events, but rather a local increase in adaptor complex recruitment into single forming CCPs.

**Figure 3.**
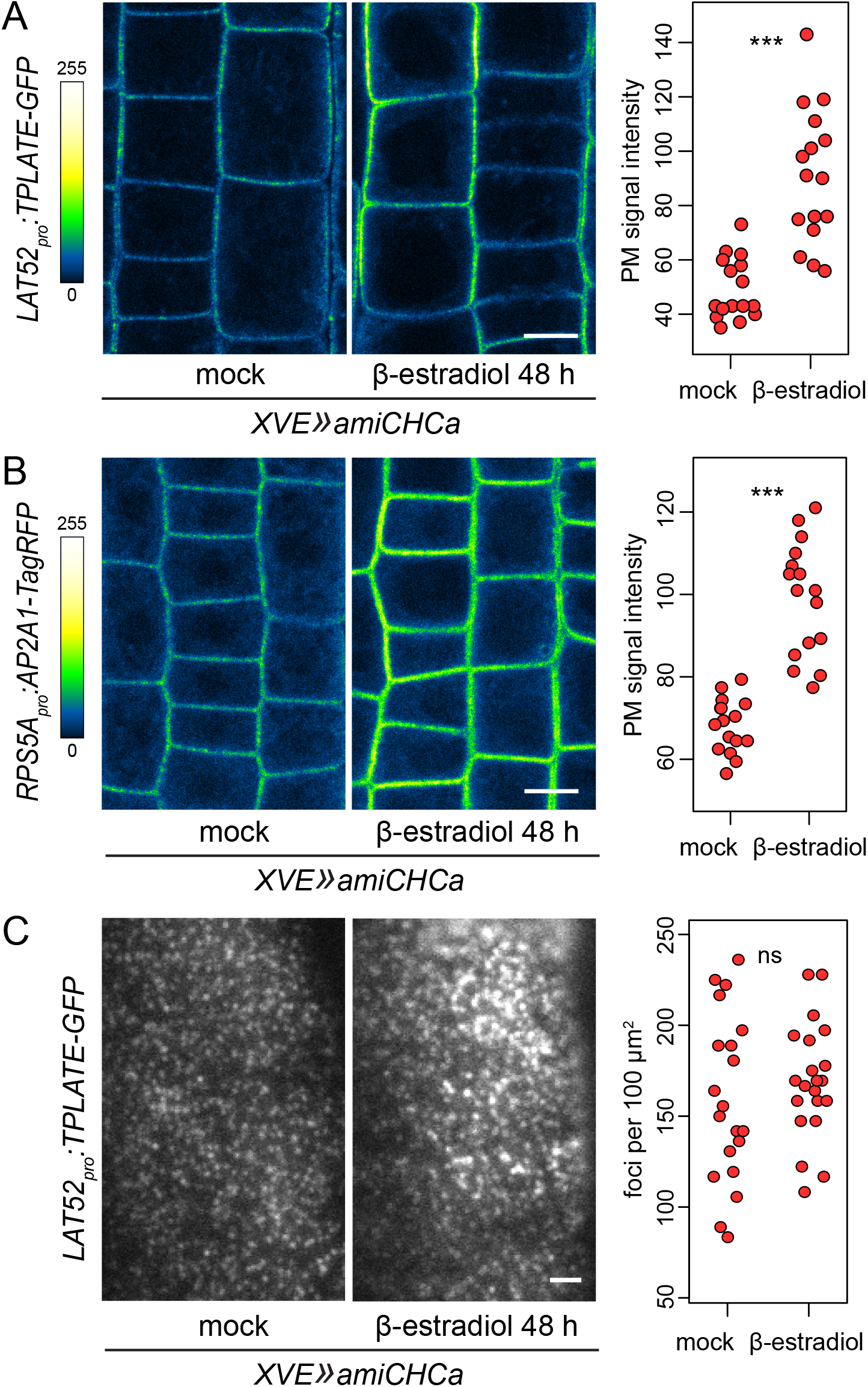
Effects of *CHC* silencing in *XVE»amiCHCa* on TPLATE-GFP and AP2A1-TagRFP. CLSM images of TPLATE-GFP (A) and AP2A1-TagRFP (B) in RAMs of control and induced seedlings of *XVE»amiCHCa*. Down-regulation of *CHC* lead to an increase in PM recruitment of the adaptor protein subunits. Graphs show quantifications of PM signal intensities from representative experiments, each data point represents a single root. TPLATE-GFP mock: 49 ± 11 a.u., n=16; β-estradiol: 90 ± 24 a.u., n=16. AP2A1-TagRFP mock: 67 ± 6 a.u., n=16; β-estradiol: 98 ± 14 a.u., n=16. Values were compared using *t* tests, *** P≤0.001. Scale bars – 10 μm. (C) TIRF images of TPLATE-GFP in roots of control and induced seedlings of *XVE»amiCHCa*. Silencing of *CHC* did not affect the density of TPLATE-positive foci at the PM. Graph shows collated quantifications of average foci density from all experiments, each data point represents a single root. Mock: 159 ± 45 foci per 100 μm^2^, n=20; β-estradiol: 169 ± 31 foci per 100 μm^2^, n=21. Values were compared using a *t* test, ns – not significant. Scale bar – 2 μm.

In summary, silencing of *CHC* in *XVE»amiCHCa* lines likely inhibited CME at a stage after the initial recruitment of adaptor protein complexes. It remains unclear why the effects on DRP1C recruitment were variable, but a decrease in average foci density also supports the conclusion.

### Effects of *CHC* silencing on internal secretory compartments

Next, we analysed the effect of *CHC* down-regulation on internal endomembrane system compartments, specifically, on GA and TGN/EE, organelles central to the activity of the secretory pathway. *XVE»amiCHCa* was transformed into lines expressing fluorescent protein fusions of GNOM-LIKE1 (GNL1), an ARF-GEF active at the GA (Teh and Moore, 2007; Richter et al., 2007); of ARFA1c, an ARF small GTPase of the ARFA1 class localized to the GA and TGN/EE compartments (Xu and Scheres, 2005; Robinson et al., 2011); and of BIG2/BEN3 (BFA-VISUALISED TRAFFICKING DEFECTIVE3), a member of the BIG ARF-GEF class acting at the TGN/EE (Kitakura et al., 2017; Richter et al., 2014). The fluorescent markers were analysed by CLSM in seedling RAM epidermis. GNL1-GFP and ARFA1c-GFP presented a normal appearance in induced *XVE»amiCHCa* lines, highlighting GA and TGN/EEs compartments of normal sizes and abundance in the cytosol (Figure 4A, B). BEN3-TagRFP too, appeared in abundant TGN/EE compartments, but the protein could be sometimes observed in slightly enlarged aggregates of high intensity signals (Figure 4C). Given that similar bright aggregations were not seen with ARFA1c-GFP, a presumed target of BIG ARF-GEFs at the TGN/EE, the effect may be specific to the BIG ARF-GEFs alone, and not indicative of a broader deficiency in the TGN/EE compartment. Taken together, the observations do not indicate major defects in the function of GA and TGN/EE compartments resulting from silencing of *CHC*.

**Figure 4.**
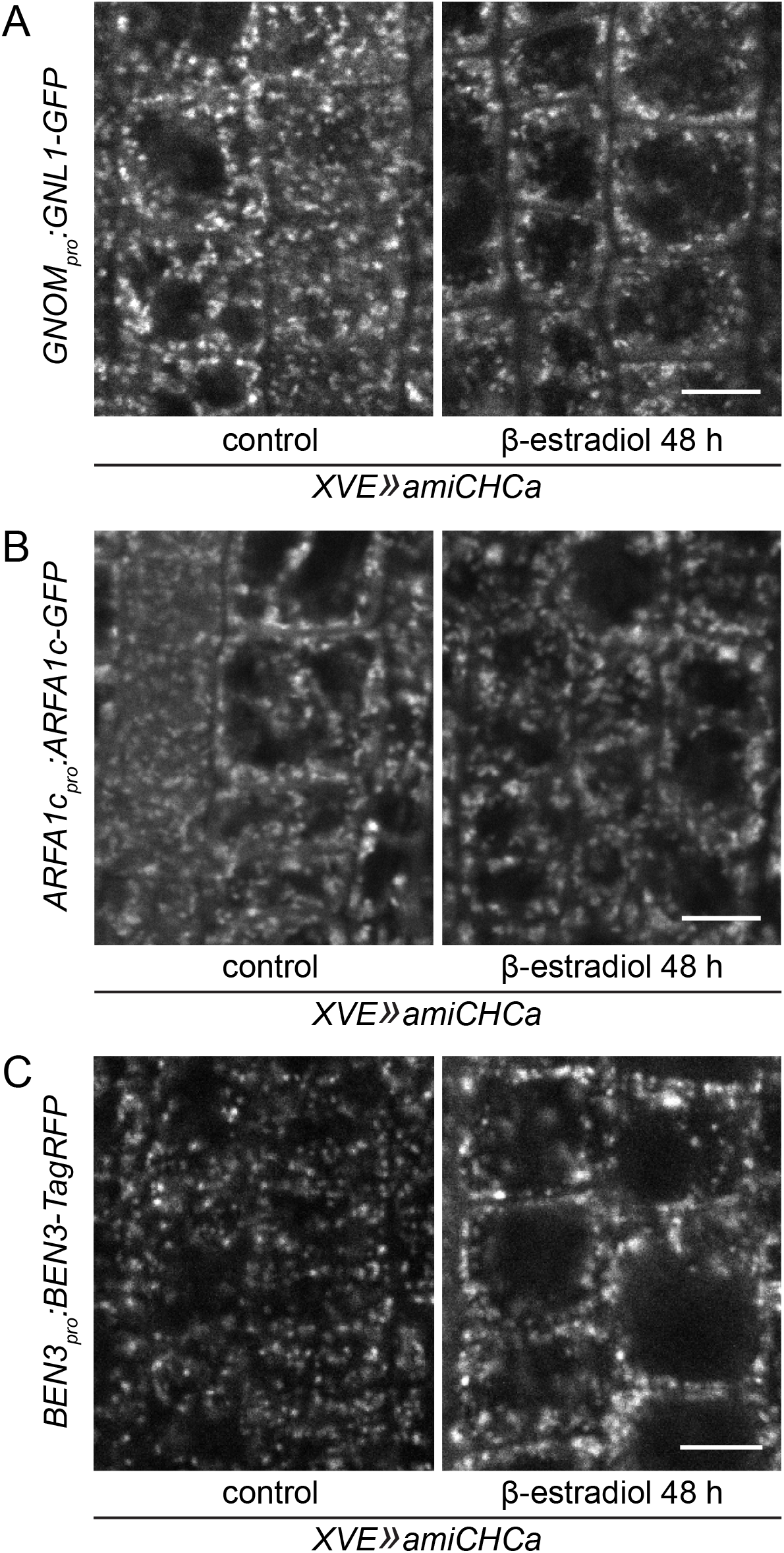
Effects of *CHC* silencing in *XVE»amiCHCa* on GNL1-GFP, ARFA1c-GFP, and BEN3-TagRFP. CLSM images of GNL1-GFP (A), ARFA1c-GFP (B), and BEN3-TagRFP (C) in RAMs of control and induced seedlings of *XVE»amiCHCa*. Following silencing of *CHC*, the distribution of GNL1-GFP and ARFA1c-GFP signals remained normal, while signals of BEN3-TagRFP occasionally appeared aggregated and brighter than in control conditions. Scale bars – 10 μm.

### Vacuolar redirection of secretory cargos as a result of *CHC* down-regulation

Finally, we characterized the effects of *CHC* down-regulation on secretion. We employed the *secRFP XVE»amiCHCa, secRFP XVE»amiCHCb*, and *secRFP XVE»amiCHCc* lines, which express an RFP variant inserted into the ER and secreted to the apoplast through the secretory pathway (Samalova et al., 2006), and additionally, introduced heat shock-inducible constructs expressing GFP fusions of PM-targeted integral membrane proteins TRANS-MEMBRANE KINASE 4 (TMK4; Dai et al., 2013; Wang et al., 2020) and PIN-FORMED1 (PIN1; reviewed in Adamowski and Friml, 2015) into *secRFP XVE»amiCHCa*. While the *HS_pro_:TMK4-GFP* construct turned out to express the TMK4-GFP fusion protein at a lower level also without a heat-shock induction (not shown), we decided nevertheless to employ this line for experiments.

When secRFP-expressing lines were investigated by CLSM in seedling RAMs after ~48 h of induction, a large portion of secRFP was located in the vacuole in all seedlings analyzed (Figure 5A), instead of being secreted to the apoplast. This observation indicates a role of clathrin in secretion. Similarly, both TMK4-GFP and PIN1-GFP were partially relocated from their normal PM localizations to the tonoplast of the main, enlarged vacuole of *XVE»amiCHC* cells, as well as, abundantly present at the membranes of smaller vacuoles often adjacent to the main vacuole (Figure 5B and 5C). Of the two integral membrane protein markers, TMK4-GFP appeared to undergo this tonoplast relocation more frequently. The abnormal localization pattern of TMK4-GFP was observed in all of 45 RAMs imaged after ~48 h of induction in the course of the experiments, essentially in all cells visible in the images, while PIN1-GFP relocated to the tonoplast in 42 of 68 RAMs imaged after ~48 h of induction, and often, only in some of the cells expressing the marker.

**Figure 5.**
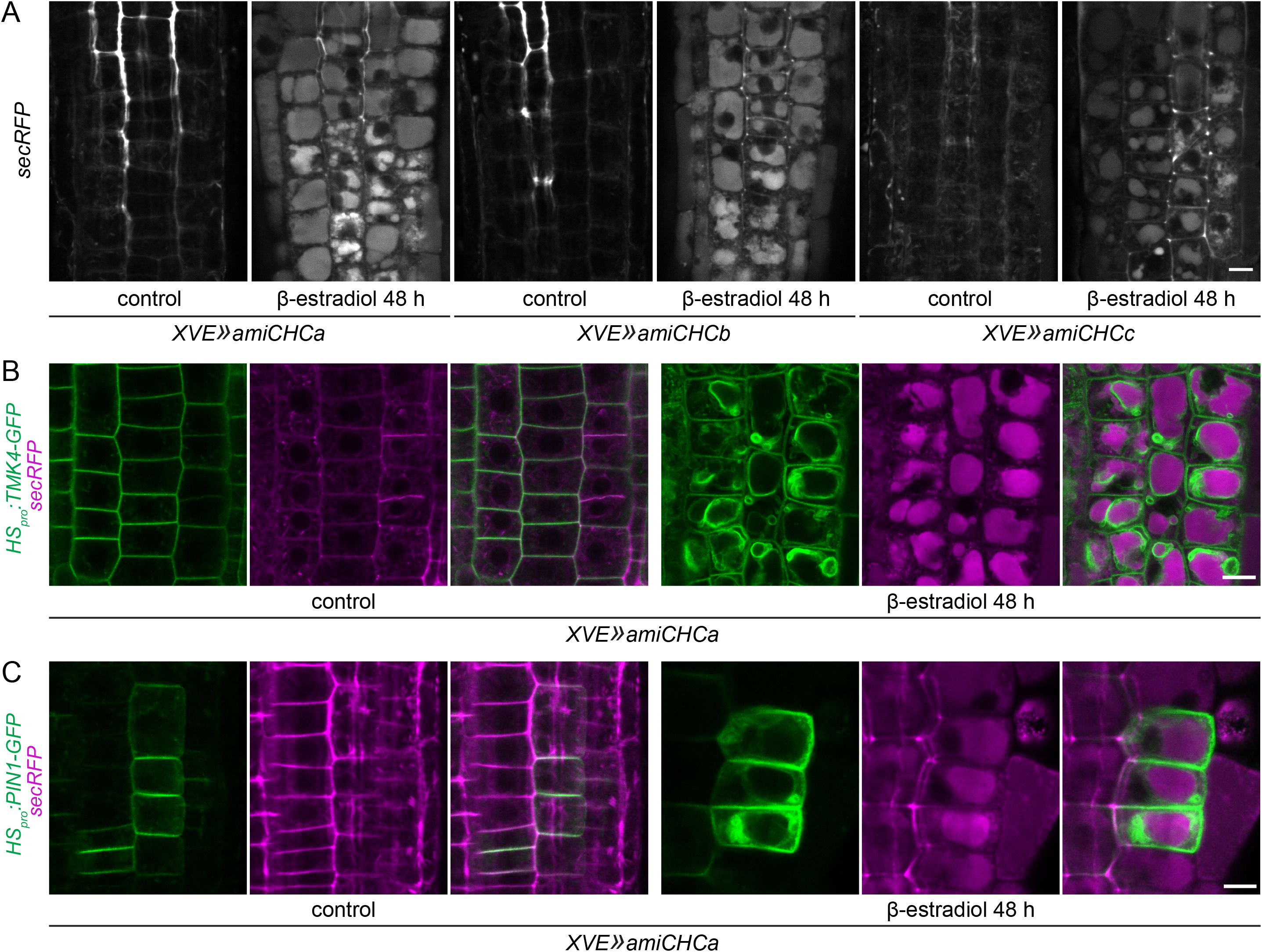
Effects of *CHC* silencing on secRFP, TMK4-GFP, and PIN1-GFP. (A) CLSM images of secRFP in RAMs of *XVE»amiCHCa, XVE»amiCHCb* and *XVE»amiCHCc* lines. Following *amiCHC* induction, large portions of secRFP are found relocated to the vacuole in all three lines. Scale bar – 10 μm. (B) and (C). CLSM colocalization images of secRFP with TMK4-GFP (B) and secRFP with PIN1-GFP (C) in RAMs of *XVE»amiCHCa* lines. Following *amiCHC* induction, TMK4-GFP and PIN1-GFP are found partially localized to the tonoplast of the central, enlarged vacuole of *XVE»amiCHCa* RAM epidermal cells and more abundantly to the tonoplasts of smaller associated vacuoles. See text for details of frequencies of the observations. Scale bars – 10 μm.

The results indicate a role of clathrin in exocytosis of soluble and transmembrane protein cargoes. Following clathrin silencing, the cargoes become relocated to the vacuole, most likely through MVBs maturating from TGNs (Scheuring et al. 2011), into which the proteins are likely non-specifically incorporated in the absence of a TGN subdomain responsible for a clathrin-dependent exocytic process. SecRFP, present in the TGN lumen, is most likely carried to the vacuolar lumen in the lumen of the MVBs, while the integral membrane proteins TMK4 and PIN1, present on the TGN membrane, are most likely carried on the outer membrane of the MVB; as new exocytic cargoes, rather than cargoes internalized from the PM for degradation, they presumably lack signals for the incorporation into intraluminal vesicles of the MVB by the ESCRT complexes (Rodriguez-Furlan et al., 2019; Cullen and Steinberg, 2018), explaining their ultimate destination at vacuolar membranes, rather than in its lumen.

It is noteworthy that following clathrin silencing, the exocytic cargoes do pass the TGN compartment, rather than being simply retained there, or at the GA. This demonstrates that the TGN as a whole remains functional, in agreement with the observations of ARFA1c-GFP and BEN3-TagRFP markers (Figure 4). Thus, down-regulation of clathrin function leads to a defect in exocytosis at the TGN compartment in a relatively specific sense. That said, the clathrin function for exocytosis at the TGN need not be entirely direct. The data should not be taken as evidence for the existence of clathrin-coated exocytic vesicles. Clathrin localized at the TGN may be contributing to exocytic activity in an indirect manner, or even, following the concepts discussed in the Introduction, exocytosis may depend on clathrin in a very indirect sense, due to a homeostatic influence of CME on an entirely clathrin-independent exocytic process. We will attempt to address these possibilities below.

### Overexpression of AUXILIN-LIKE1 inhibits CME preferentially over clathrin-dependent exocytosis

Overexpression of putative uncoating factors for endocytic CCVs AUXILIN-LIKE1/2 (Adamowski et al., 2018) leads to an inhibition of CME, as indicated by three lines of evidence: (1) inhibition of endocytosis of the FM4-64 endocytic tracer and of protein cargoes, (2) changes in the PM recruitment and dynamics of proteins participating in CME, and (3) an excess accumulation of membrane material at the cell periphery. This last observation is interpreted as resulting from a disruption of the normal balance between endo- and exocytosis trafficking, where continued exocytosis in the absence of endocytosis gradually leads to a build-up of membrane at the cell periphery. The effect of AUXILIN-LIKE1/2 overexpression on the secretory pathway was not explicitly addressed. Given the finding that silencing of *CHC* does affect exocytosis, we assessed exocytosis in *XVE»AUXILIN-LIKE1* by introducing *secRFP, HS_pro_:TMK4-GFP*, and *HS_pro_:PIN1-GFP* transgenes into this line.

First, we generated a cross between *XVE»AUXILIN-LIKE1* and *secRFP*. Regrettably, the *XVE»AUXILIN-LIKE1* transgene became partially silenced, and the overexpression phenotypes were weak in comparison to the original line, as evidenced by the ability of seeds to germinate and develop on media supplemented with *β*-estradiol (not shown), contrasted with a typically observed germination arrest (Adamowski et al., 2018). While in the original *XVE»AUXILIN-LIKE1/2* lines, the overexpression phenotypes were studied after 1 d of induction, in this case we carried out a prolonged, 2 d long induction on *β*-estradiol. Having screened multiple seedlings, we found few exhibiting clear AUXILIN-LIKE1 overexpression phenotypes, matching in their degree the phenotypes typical for 2 d long induction in the reference lines. In these seedlings, secRFP could be seen localized to the vacuole (Figure 6A). The observation indicates that overexpression of AUXILIN-LIKE1, like silencing of *CHC*, affects exocytosis. A reliable quantitative comparison of the effect with *XVE»amiCHC* lines was not possible not due to the rare occurrence of seedlings with correct phenotypes, and due to an additional silencing of secRFP expression in this cross compared with *secRFP XVE»amiCHC* lines (not shown).

**Figure 6.**
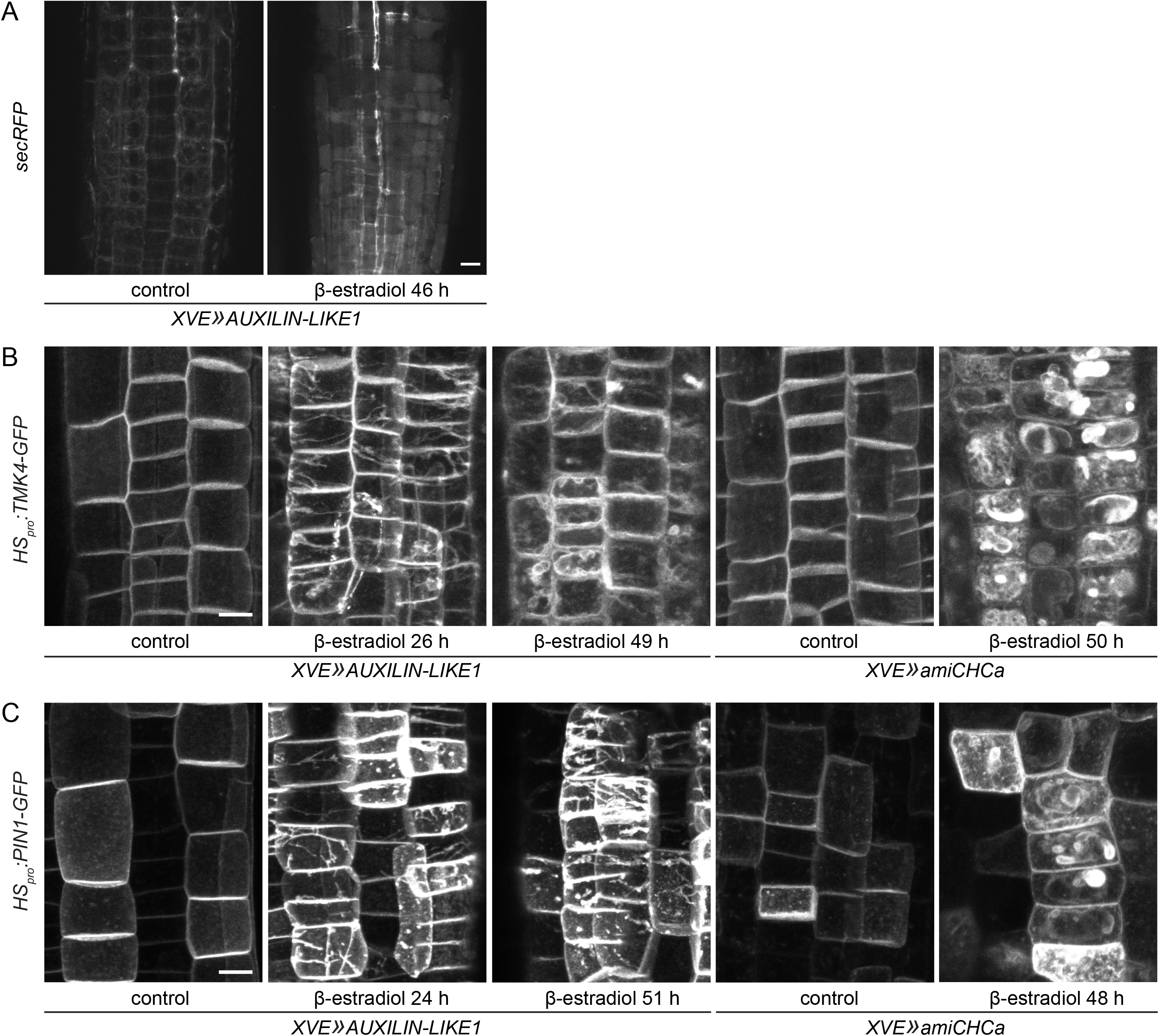
Effects of AUXILIN-LIKE1 overexpression on secRFP, TMK4-GFP, and PIN1-GFP. (A) CLSM images of secRFP in RAMs of *XVE»AUXILIN-LIKE1* line. In seedlings exhibiting AUXILIN-LIKE1 overexpression phenotypes in this cross, secRFP is found partially localized to the vacuole after 46h of induction. Scale bar – 10 μm. (B) and (C). Maximum projections of 9 μm z-stacks of CLSM images of TMK4-GFP (B) and PIN1-GFP (C) in RAMs of *XVE»AUXILIN-LIKE1* and *XVE»amiCHCa* taken after indicated times of induction. Overexpression of AUXILIN-LIKE1 for 24-26 h typically leads to deposition of excess filamentous membrane material at the PM, where TMK4-GFP and PIN1-GFP are found. TMK4-GFP, but not PIN1-GFP, is typically found localized to the tonoplast after 49-51 h induction. Filamentous membrane depositions are typically not observed in *XVE»amiCHCa* lines induced for 48-50 h. See text for details of frequencies of the observations and Suppl. Figure 3A and B for exceptions. Scale bars – 10 μm.

For a more precise analysis of exocytic activity following AUXILIN-LIKE1 overexpression we turned to *XVE»AUXILIN-LIKE1* lines transformed with *HS_pro_:TMK4-GFP* and *HS_pro_:PIN1-GFP* constructs. In these lines, the AUXILIN-LIKE1 overexpression phenotypes were normal, providing for reliable and well-controlled experimentation. As in the case of *secRFP XVE»amiCHCa HS_pro_:TMK4-GFP* lines, TMK4-GFP expression was present also without heat shock induction (not shown). We first tested the effect of AUXILIN-LIKE1 overexpression on TMK4-GFP and PIN1-GFP exocytosis after 24-26 h of AUXILIN-LIKE1 expression induction. At this point, the consequences of AUXILIN-LIKE1 overexpression on CME are well visible (Adamowski et al., 2018) and in the analysed lines, they too were evident by the abnormal morphology of the RAMs. Yet, in none of the 30 and 49 RAMs expressing TMK4-GFP and PIN1-GFP, respectively, were the exocytic cargoes relocated to the vacuole, indicating that the exocytic pathway functioned normally at this stage. Instead, in a majority of RAMs (TMK4-GFP: 20 of 30 and PIN1-GFP: 33 of 49), the PM pool of each protein was distributed in a form of thick filaments in at least part of the epidermal cells (Figure 6B and 6C), representing the previously discussed excess accumulation of membrane material characteristic for *XVE»AUXILIN-LIKE1/2* lines. Note that these filaments are an accumulation of the membrane itself, rather than an accumulation of an overexpressed fluorescent protein fusion, as they were observed with the FM4-64 membrane stain in a line not expressing any cargo reporter (Adamowski et al., 2018).

When assessed at a later time point of 49-51 h of AUXILIN-LIKE1 expression induction, TMK4-GFP relocated to the tonoplast in 9 of 11 RAMs (Figure 6B), in some cases co-existing with a localization in the PM filaments. In the case of PIN1-GFP, localization to the tonoplast was seen in only 4 of 44 RAMs (Suppl. Figure 3A) while deposition in PM-localized filaments, similar to the earlier times of induction, in 25 of 44 RAMs (Figure 6C). This rarer tendency of PIN1-GFP for relocation to the tonoplast, compared with TMK4-GFP, reflects a similar difference in the apparent sensitivity of the two markers to a disturbance of clathrin function in *XVE»amiCHCa*, described in the previous section.

For the purpose of a reliable comparison between *AUXILIN-LIKE1*-overexpressing and *CHC-*silencing lines, a part of the CLSM experiments with *XVE»amiCHCa HS_pro_:TMK4-GFP* and *XVE»amiCHCa HS_pro_:PIN1-GFP* described earlier, was performed with the use of z-stacks, capturing both the PM surface and the cell interior of the epidermal cells (Figure 6B and 6C, right), similarly to the experiments on *XVE»AUXILIN-LIKE1* lines. By this, we systematically controlled for the possible formation of filamentous membrane deposits in *XVE»amiCHCa*. Structures similar to those observed in *XVE»AUXILIN-LIKE1* were found in only 2 of 46 RAMs captured as z-stacks in *XVE»amiCHCa HS_pro_:PIN1-GFP* lines induced for ~48 h (Suppl. Figure 3B), and in none of 27 RAMs of *XVE»amiCHCa HS_pro_:TMK4-GFP* captured in similar conditions, even though a portion of each protein was always still delivered to the PM. Given that the formation of such structures precedes tonoplast relocation of the exocytic cargoes in *XVE»AUXILIN-LIKE1*, we also tested *XVE»amiCHCa* at an earlier stage of ~31 h induction, but found only cases of milder tonoplast relocation of TMK4-GFP and PIN1-GFP (Suppl. Figure 3C and 3D) and no tendency for excess membrane depositions at the PM.

Taken together, in contrast to *XVE»amiCHCa*, overexpression of AUXILIN-LIKE1, at least at earlier points of expression induction, inhibits endocytosis preferentially, while not having an equal effect on exocytosis. This is manifested by an excess deposition of membranes at the cell periphery without a concomitant relocation of cargoes to the tonoplast. With regard to the question of potential homeostatic regulations between endo- and exocytosis, it is highly unlikely that CME, through a homeostatic mechanism, is tied to the rates of secretion, as in this case, overexpression of AUXILIN-LIKE1/2 would need to present phenotypes not different from those resulting from silencing of *CHC*. Returning to the role of clathrin in exocytosis, the comparison indicates that the impact of *CHC* silencing on exocytosis likely manifests a direct function of TGN-localized clathrin in exocytosis, instead of an indirect, homeostatic reaction of the exocytic pathway to the inhibition of clathrin function in CME.

### Assessment of the rates of CME in secretory mutants

To explore further the possible homeostatic regulations between endo- and exocytosis, we asked the complementary question of whether endocytic rates are reactive to the cellular rates of secretion. We addressed this through an assessment of CME following interference with exocytosis. We crossed CLC2 and TPLATE fluorescent reporter lines into selected mutants deficient in secretion and measured the rates of CME using TIRF microscopy. We employed *ap1m2-1*, a mutant of the μ subunit of the TGN-localized AP-1 clathrin adaptor complex, deficient both in secretion and vacuolar traffic (Park et al., 2013; see Introduction); *echidna* (*ech*), a mutant of a TGN-localized secretory component ECH likely associated with the Rab and ARF small GTPase machinery (Gendre et al., 2011, 2013; Jonsson et al., 2017; Ravikumar et al., 2018); and *sec5a-2 sec5b-1*, a mutant in the SEC5 component of the exocyst, a tethering complex responsible for the docking of SVs to the PM in final steps of exocytosis (Zhu et al., 2018; Žárský et al., 2013). All these mutants exhibit significant morphological deficiencies characterized by a clear delay in growth and development in seedlings after 7 d of culture, when TIRF imaging was conducted. We evaluated the state of CME in the mutants by quantifying the density of CCPs marked by CLC2-GFP and by TPLATE-GFP in the early elongation zone of the mutant seedling roots.

First, we evaluated the density of CLC2-GFP signals in the mutant lines. In a total sample of 30 to 45 cells taken from 18 to 20 seedlings of each genotype in the course of the experiments, we found a slight but significant reduction in the average density of CLC2-GFP-positive foci at the PMs of all three secretory mutants (Figure 7A). To ascertain the validity of these results, we repeated the statistical tests after removing two outliers of very high values from the wild type sample and one outlier of a very high value from *sec5a-2 sec5b-1*. Following this, the average densities of CLC2-GFP-positive foci in all three mutants were still significantly lower than in the wild type sample (Suppl. Figure 4). Next, we conducted similar experiments with the TPLATE-GFP marker protein. In a total sample of 48 to 91 cells taken from 18 to 34 seedlings of each genotype, we found only the *ap1m2-1* mutant to have a significant reduction in the density of TPLATE-GFP foci density, while *ech* and *sec5a-2 sec5b-1* had foci density not significantly different from the control.

**Figure 7.**
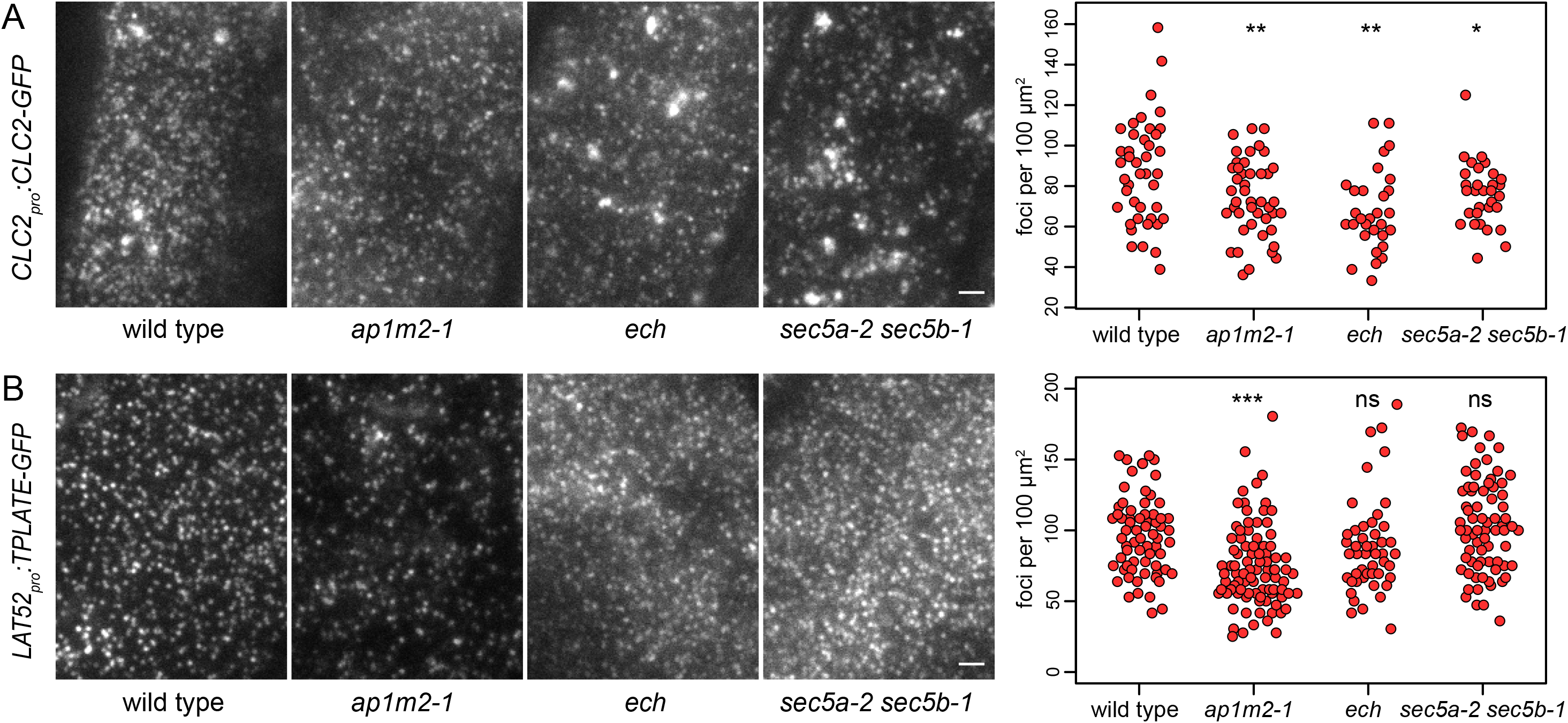
Rates of CME in secretory mutants. Representative TIRF microscopic images and graphs visualising foci densities of CME markers CLC2-GFP (A) and TPLATE-GFP (B) in seedling roots of *ap1m2-1, ech*, and *sec5a-2 sec5b-1* mutants. Scale bars – 2 μm. Graphs show collated data from all experiments, each data point represents a measurement from a single cell. CLC2-GFP wild type 87 ± 25 foci per 100 μm^2^, n=42; *ap1m2-1* 74 ± 19 foci per 100 μm^2^, n=45; *ech* 68 ± 20 foci per 100 μm^2^, n=30; *sec5a-2 sec5b-1* 76 ± 15 foci per 100 μm^2^, n=32. TPLATE-GFP wild type 97 ± 27 foci per 100 μm^2^, n=68; *ap1m2-1* 75 ± 29 foci per 100 μm^2^, n=91; *ech* 88 ± 33 foci per 100 μm^2^, n=48; *sec5a-2 sec5b-1* 103 ± 33 foci per 100 μm^2^, n=73. Mutant values were compared with the wild type using *t* tests, * P≤0.05, ** P≤0.01, *** P≤0.001, ns – not significant. See Suppl. Fig. 4 for a statistical evaluation of CLC2-GFP foci density after removal of outliers of high value.

Taken together, we found support for the conclusion that the rate of CME is, to some degree, reduced in the *ap1m2-1* mutant, but we did not find consistent evidence for a reduction of rates of CME in *ech* and *sec5a-2 sec5b-1* mutants. Thus, our results do not provide a strong support for a homeostatic adjustment of the rates of endocytosis to the rates of exocytosis.

## Discussion

In this work, we addressed two related questions about the function of the plant cell’s endomembrane system: that of the activity of TGN-localized clathrin, and that of potential regulatory mechanisms mutually adjusting rates of endocytosis and exocytosis as a part of a homeostatic control component within the endomembrane system.

### An involvement of clathrin in exocytosis

The role of clathrin at the TGN has been a matter of much debate. Indirect evidence, especially obtained with the use of mutants in components associated with clathrin in the vesicle formation process at the TGN, argue for the involvement of CCVs either in vacuolar, or in exocytic, trafficking pathways (Park et al., 2013; Wang et al., 2014; Robinson and Pimpl, 2014; Feraru et al., 2010; Zwiewka et al., 2011; Sauer et al., 2013; Heinze et al., 2020; Collins et al., 2020). Our results from inducible downregulation of *CHC* expression support a function of clathrin in exocytosis of soluble and integral membrane protein cargo. The observations from *amiCHC* lines are specific to the degree that exocytic cargoes were redirected from the TGN towards the vacuole, likely in maturating MVBs, indicating that a clathrin-dependent exocytic process at the TGN was disrupted specifically, without an associated major defect of the organelle, which would wholly arrest the function of this compartment. That said, the nature of the experiments presented does not provide evidence for a fully direct role of clathrin in exocytosis, that is, for the existence of clathrin-coated vesicles forming at the TGN compartment and destined for fusion with the PM. Clathrin at the TGN compartment m may support exocytosis indirectly. For instance, clathrin at the TGN may be involved in a process of sorting proteins out of a TGN domain destined to maturate into MVBs, and into domains destined to be fragmented into canonical SVs. However, this proposition would be at odds with the perception of the secretory route to the PM as a default pathway, and could, at best, explain sorting of integral membrane proteins, but not of secRFP, due to a lack of sorting signals for segregation into CCPs in this protein. In summary, while not providing mechanistic details of the function of clathrin in exocytosis, our *CHC* loss of function data are an interesting addition to the discussion about the function of TGN-localized clathrin in plant cells.

### Homeostatic control of exocytosis and endocytosis

Theoretical considerations, as well as some experimental results (Riezman, 1985; Steer, 1988; Larson et al., 2017, Yan et al., 2021), suggest the existence of mechanisms within the endomembrane system that adjust the rates of individual trafficking processes to one another on a cellular scale. We scrutinized whether such homeostatic controls may exist between the rates of major trafficking routes in the cell: endo- and exocytosis. Through a comparison of *amiCHC* lines with lines overexpressing a putative uncoating factor for endocytic CCVs, AUXILIN-LIKE1, where inhibition of endocytosis preceded any visible interference with exocytic traffic of integral membrane cargoes, we conclude that a homeostatic mechanism modulating exocytosis to the current rates of endocytosis is unlikely to be present in plant cells. This conclusion appears intuitive when secretion is perceived as a fundamental cellular activity, required for functions as essential as the construction of the cell wall. Still, experiments on secretory activity in loss-of-functions mutants of components involved solely in CME, or on similar lines down-regulating their activity, could be a valuable future addition to the data presented here with the use of *XVE»AUXILIN-LIKE1* line.

A complementary assessment of a potential mechanism adjusting rates of endocytosis to the rates of secretion, by direct measurements of CME in secretory mutants, yielded variable results. The *ap1m2-1* mutant, in which a reduction in rates of CME was detected with both CLC2-GFP and TPLATE-GFP markers, may be deficient in secretion to a higher degree than the other mutants analysed, explaining why a decrease in CME was also more reliably detected. It may be alternatively proposed that loss of AP-1 function affects an exocytic pathway distinct from that involving ECH and/or the exocyst, and that this pathway only, is mechanistically tied with the rates of CME. Finally, it could be proposed that a hypothetical control mechanism acts on clathrin recruitment to forming CCPs, but not on the earlier TPLATE complex recruitment to the PM, explaining why a reduction of CLC2-GFP foci density is observed more readily than a reduction of TPLATE-GFP foci density in secretory mutants. Currently, these possibilities remain unexplored, and when all the gathered data are considered together, we tentatively conclude that our results do not clearly support the existence of a generally supposed (Steer, 1988; Larson et al. 2017, Yan et al., 2021) homeostatic mechanism adjusting rates of CME to the status of the secretory pathway.

## Materials and Methods

### Plant material

The following previously described *A. thaliana* lines were used in this study: *secRFP* (Samalova et al., 2006; NASC ID N799370), *CLC2_pro_:CLC2-GFP* (Konopka et al., 2008), *CLC2_pro_:CLC2-GFP UBQ10_pro_:mCh-AUXILIN-LIKE1* (Adamowski et al., 2018), *LAT52_pro_:TPLATE-GFP RPS5A_pro_:AP2A1-TagRFP tplate* (Gadeyne et al., 2014), *DRP1C_pro_:DRP1C-GFP* (Konopka et al., 2008)*, GNOM_pro_:GNOM-LIKE1-GFP* (Adamowski et al., 2021), *ARFA1c_pro_:ARFA1c-GFP* (Xu and Scheres, 2005), *BEN3_pro_:BEN3-TagRFP* (Kitakura et al., 2017), *XVE»AUXILIN-LIKE1* (Adamowski et al., 2018), *ap1m2-1* (Park et al., 2013), *echidna* (Gendre et al., 2001), *sec5a-2 sec5b-1* (Zhu et al., 2018). Lines generated as part of this study are listed in Suppl. Table 1 and primers used for genotyping in Suppl. Table 2.

### *In vitro* cultures of *Arabidopsis* seedlings

Seedlings were grown in *in vitro* cultures on half-strength Murashige and Skoog (½MS) medium of pH=5.9 supplemented with 1% (w/v) sucrose and 0.8% (w/v) phytoagar at 21 °C in 16h light/8h dark cycles with Philips GreenPower LED as light source, using deep red (660nm)/far red (720nm)/blue (455nm) combination, with a photon density of about 140μmol/(m^2^s) +/- 20%. Beta-estradiol (Sigma-Aldrich) was solubilized in 100% ethanol to 5 mg/mL stock concentration and added to ½MS media during preparation of solid media to a final concentration of 2.5 μg/mL. Seedlings of *XVE»amiCHC* and *XVE»AUXILIN-LIKE1* lines were induced by transferring to beta-estradiol-supplemented media at day 3 or 4. Heat shock induction of *HS_pro_:TMK4-GFP* and *HS_pro_:PIN1-GFP* was initiated 7-8 h before live imaging and carried out for 1 h at 37 °C.

### Light microscopy

Images of *XVE»amiCHC* seedlings developing in *in vitro* cultures were taken with Leica EZ4 HD stereomicroscope, and microscopic images of RAMs were taken with Olympus BX53 light microscope.

### Confocal Laser Scanning Microscopy

4 to 5 d old seedlings were used for live imaging with Zeiss LSM800 confocal laser scanning microscope with 20X lens. For comparative studies of TMK4-GFP and PIN1-GFP localizations in *XVE»amiCHCa* and *XVE»AUXILIN-LIKE1* lines, z-stacks of 9 images at 1 μm intervals were captured. Bleaching of CLC2-GFP signals was performed by exposing the sample to high laser for around 10 seconds. Measurements of PM signal intensities of TPLATE-GFP and AP2A1-TagRFP were performed in Fiji (https://imagej.net/Fiji) as mean grey values of a line of 5 pixel width drawn over multiple PMs in each CLSM image.

### Total Internal Reflection Fluorescence microscopy

Early elongation zone of roots in excised ~1 cm long root tip fragments from 7d old seedlings were used for TIRF imaging. Imaging was performed with Olympus IX83 TIRF microscope, using a 100X TIRF lens with an additional 1.6X magnification lens in the optical path. Time lapses of 100 frames at 1 s intervals with exposure times of 200 ms, or single snapshots of 200 ms exposure, were taken, depending on the experiment. DRP1C-GFP, CLC2-GFP and TPLATE-GFP foci were counted in square regions of 36 μm^2^ taken from the captured TIRF images or movies.

### Molecular cloning

All constructs generated in this study are listed in Suppl. Table 3 and primers used for cloning in Suppl. Table 2. Artificial microRNA sequences targeting *CHC* genes were designed using Web MicroRNA Designer website (http://wmd3.weigelworld.org/). The following amiRNA sequences were employed: amiCHCa TCTCGCAGTACTTACCCACAA; amiCHCb TGCAAAATTTACTGCTCCCTG; amiCHCc TTATTTGACCAGTCTTGGCAG. amiRNA constructs were cloned by overlap extension PCR with iProof High Fidelity polymerase (BioRad), following a protocol published on the Web MicroRNA Designer website. The amiRNA sequences were introduced into pDONR221 Gateway entry vectors (Invitrogen) by BP Clonase, and subsequently combined by LR Clonase II with UBQ10-promoter driven XVE elements from p1R4-pUBQ10:XVE (Siligato et al., 2016) in pH7m24GW,3, pK7m24GW,3 (*amiCHCa*) and pB7m24GW,3 (*amiCHCa,b,c*) expression vectors (Karimi et al., 2002). TMK4 and PIN1-GFP-2 coding sequences were cloned into pDONR221 Gateway entry vectors by BP Clonase. Heat-shock inducible expression vectors were generated by LR Clonase II with proHS/pDONRP4P1r and GFP/pDONRP2rP3 (for TMK4-GFP) in expression vectors pK7m34GW (TMK4-GFP) and pK7m24GW,3 (PIN1-GFP-2).

### Quantitative reverse transcriptase PCR

Total RNA was isolated from 5d old seedlings using RNeasy Plant Mini Kit (Qiagen). cDNA was synthetised using iScript™ cDNA Synthesis Kit (Bio-Rad). qPCR was performed with Luna^®^ reagent (New England Biolabs). 2 primer pairs detecting both *CHC1* and *CHC2* transcripts were employed. *TUB2* and *PP2AA3* were used as reference genes. Primer sequences are listed in Suppl. Table 2.

### Accession numbers

Sequence data from this article can be found in the GenBank/EMBL libraries under the following accession numbers: CHC1 (AT3G11130), CHC2 (AT3G08530), CLC2 (AT2G40060), TPLATE (AT3G01780), AP2A1 (AT5G22770), DRP1C (AT1G14830), GNOM-LIKE1 (AT5G39500), ARFA1c (AT2G47170), BEN3/BIG2 (AT3G60860), TMK4 (AT2G42330), PIN1 (AT1G73590), AUXILIN-LIKE1 (AT4G12780), AP1M2 (AT1G60780), ECHIDNA (AT1G09330), SEC5A (AT1G76850), SEC5B (AT1G21170), TUB2 (AT5G62690), PP2AA3 (AT1G13320).

## Supplemental Data files

Suppl. Figure 1. Morphology of *XVE»amiCHC* lines on control media

Suppl. Figure 2. qPCR analysis of *secRFP XVE»amiCHC* lines

Suppl. Figure 3. Additional data on TMK4-GFP and PIN1-GFP localizations in *XVE»AUXILIN-LIKE1* and *XVE»amiCHCa* lines

Suppl. Figure 4. Statistical evaluation of CLC2-GFP foci density after removal of outliers

Suppl. Movie 1. TIRF time lapse of CLC2-GFP in control conditions

Suppl. Movie 2. TIRF time lapse of CLC2-GFP in *XVE»amiCHCa* induced for 45 h

Suppl. Table 1. Lines generated as part of this study

Suppl. Table 2. Primers used in this study

Suppl. Table 3. Constructs generated as part of this study

## Author Contributions

M.A. and J.F. designed research, analyzed data and wrote the manuscript. M.A. and I.M. performed research.

## Acknowledgements

The authors wish to acknowledge Dr. Paweł Baster for cloning PIN1-GFP-2/pDONR221, Prof. Ari Pekka Mähönen for sharing the p1R4-pUBQ10:XVE plasmid, and Prof. Ying Gu for sharing seeds of the *sec5* mutant. M.A. would like to thank Dr. Xixi Zhang and Prof. Sebastian Bednarek for inspiring discussions. This work was supported by the Austrian Science Fund (FWF): I 3630-B25.

---

**Suppl. Figure 1. Morphology of *XVE»amiCHC* lines on control media**

(A) *secRFP XVE»amiCHC* lines grown on a control medium for 6 d.

(B) Microscopic images of RAMs of *secRFP XVE»amiCHC* lines from control media. Scale bar – 10 μm.

**Suppl. Figure 2. qPCR analysis of *secRFP XVE»amiCHC* lines**

The lines were induced on β-estradiol for 2 days before RNA isolation. Transcript levels of *CHC1* and *CHC2* were assessed with two primer pairs, each recognizing transcripts from both homologous genes, against reference genes *TUB2* and *PP2AA3*. Graphs represent mean ± SD from two biological replicates. Due to a technical error, 1 biological replicate was assessed for mock-treated *secRFP XVE»amiCHCc*.

**Suppl. Figure 3. Additional data on TMK4-GFP and PIN1-GFP localizations in *XVE»AUXILIN-LIKE1* and *XVE»amiCHCa* lines**

All panels are maximum projections of 9 μm z-stacks of CLSM images. See text for details of frequencies of the observations. Scale bars – 10 μm.

(A) Localization of PIN1-GFP to the vacuole following AUXILIN-LIKE1 overexpression for 51 h.

(B) Presence of filamentous membrane depositions highlighted with PIN1-GFP in *XVE»amiCHCa* line induced for 48 h.

(C) and (D). Mild re-localization of TMK4-GFP (C) and PIN1-GFP (D) to the tonoplast after 31 h of *amiCHCa* induction.

**Suppl. Figure 4. Statistical evaluation of CLC2-GFP foci density after removal of outliers**

Graph visualising foci density of CLC2-GFP in seedling roots of *ap1m2-1, ech*, and *sec5a-2 sec5b-1* mutants. Each data point represents a measurement from one cell. Data points in grey are outliers of high values removed from this analysis. Mutant values were compared with the wild type control using *t* tests after removal of outliers, * P≤0.05, ** P≤0.01.

