## Supplemental Figures and Tables for "The role of clathrin in exocytosis and the mutual regulation of endo- and exocytosis in plant cells"

Suppl. Figure 1

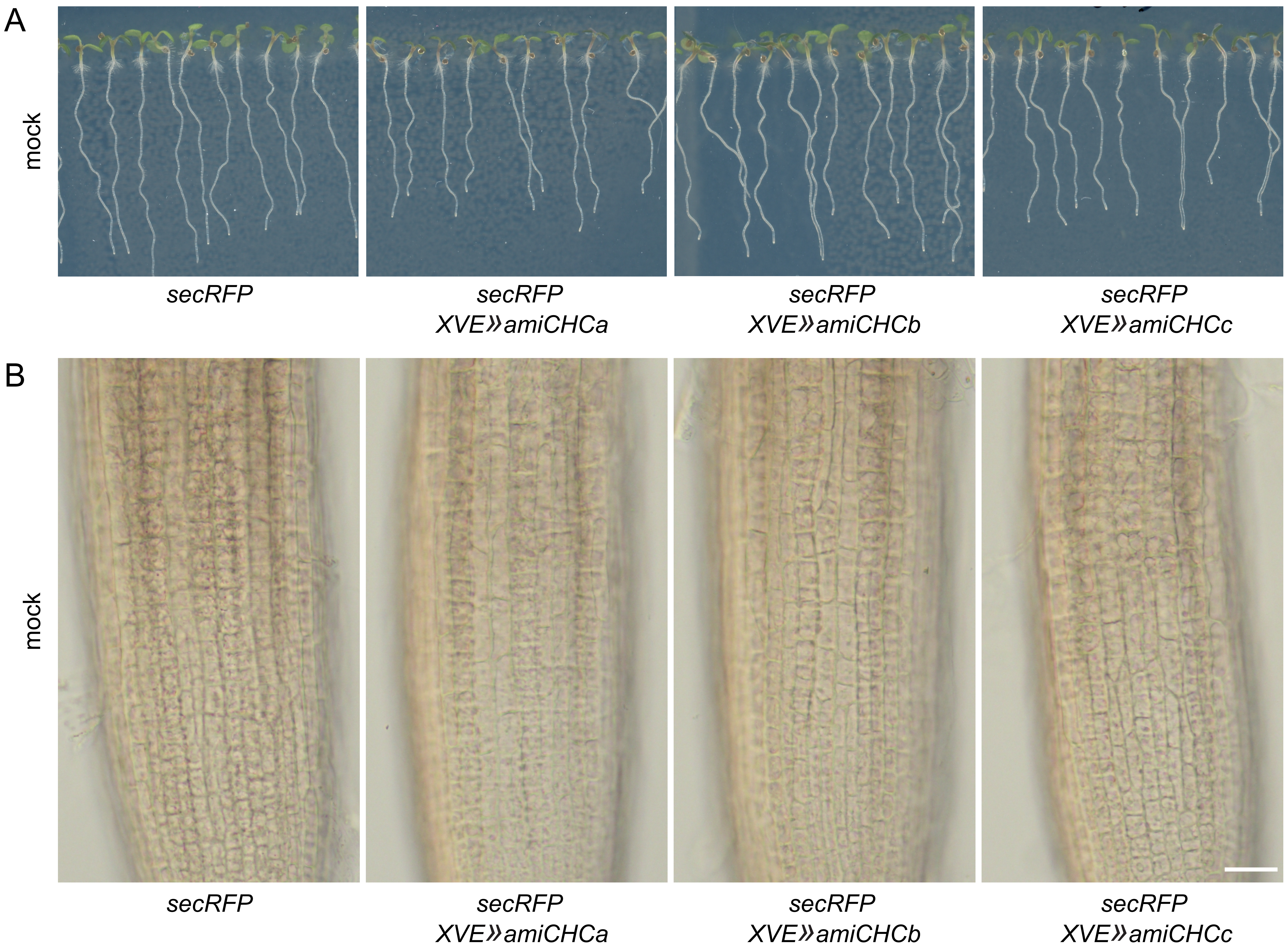


**Suppl. Figure 1. Morphology of XVE»amiCHC lines on control media**
Related to Figure 1.
(A) secRFP XVE»amiCHC lines grown on a control medium for 6 d.
(B) Microscopic images of RAMs of secRFP XVE»amiCHC lines from control media. Scale bar – 10 µm.

### Suppl. Figure 2

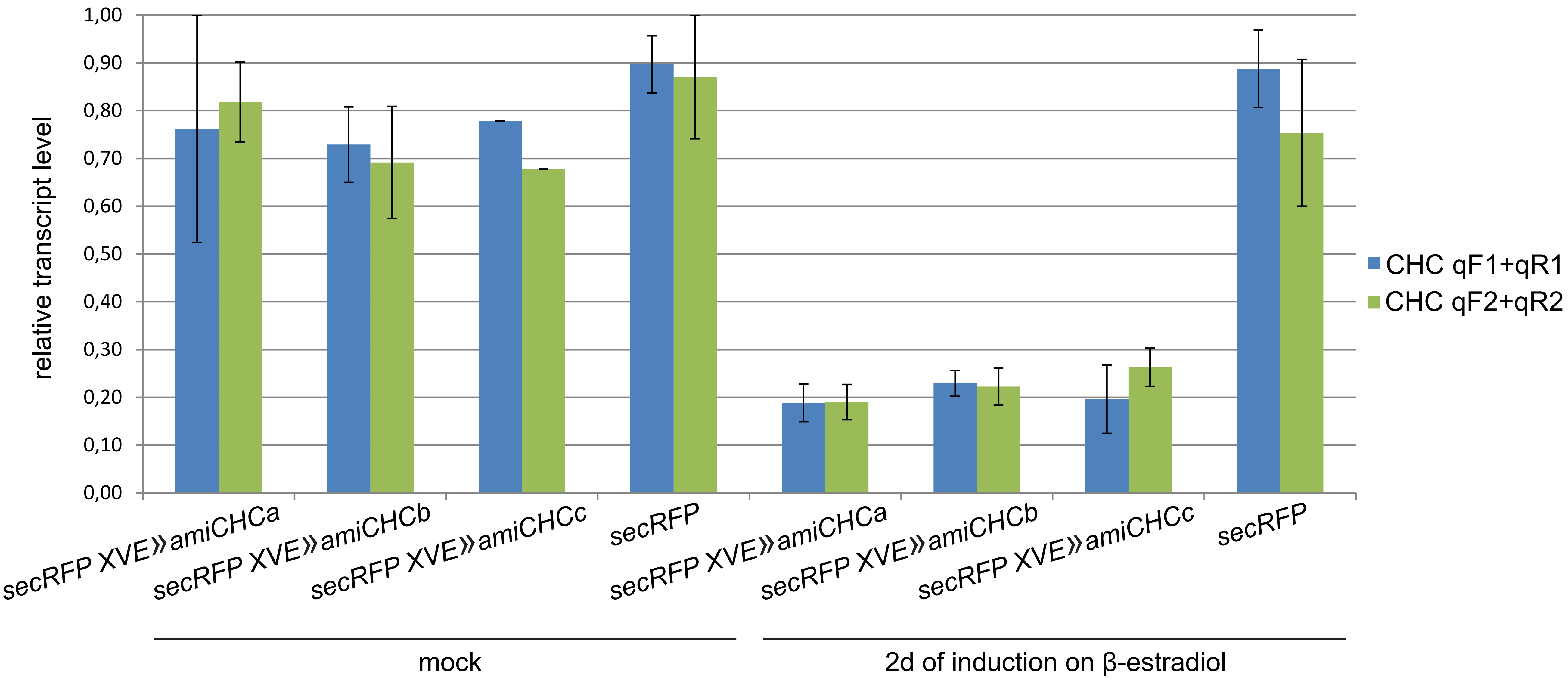


**Suppl. Figure 2. qPCR analysis of secRFP XVE»amiCHC lines**
Related to Figure 1.
The lines were induced on β-estradiol for 2 days before RNA isolation. Transcript levels of CHC1 and CHC2 were assessed with two primer pairs, each recognizing transcripts from both homologous genes, against reference genes TUB2 and PP2AA3. Graphs represent mean ± SD from two biological replicates. Due to a technical error, 1 biological replicate was assessed for mock-treated secRFP XVE»amiCHCc.

### Suppl. Figure 3

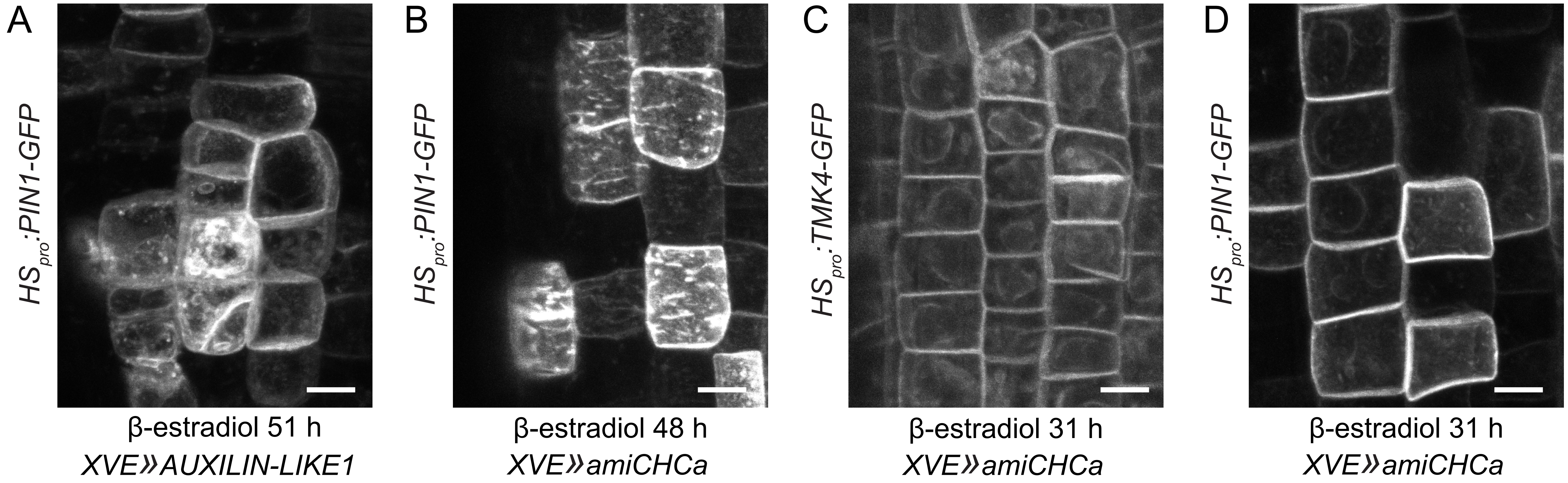


**Suppl. Figure 3. Additional data on TMK4-GFP and PIN1-GFP localizations in XVE»AUXILIN-LIKE1 and XVE»amiCHCa lines**
Related to Figure 6.
All panels are maximum projections of 9 µm z-stacks of CLSM images. See text for details of frequencies of the observations.
Scale bars – 10 µm.
(A) Localization of PIN1-GFP to the vacuole following AUXILIN-LIKE1 overexpression for 51 h.
(B) Presence of filamentous membrane depositions highlighted with PIN1-GFP in XVE»amiCHCa line induced for 48 h.
(C) and (D). Mild re-localization of TMK4-GFP (C) and PIN1-GFP (D) to the tonoplast after 31 h of amiCHCa induction.

Suppl. Figure 4

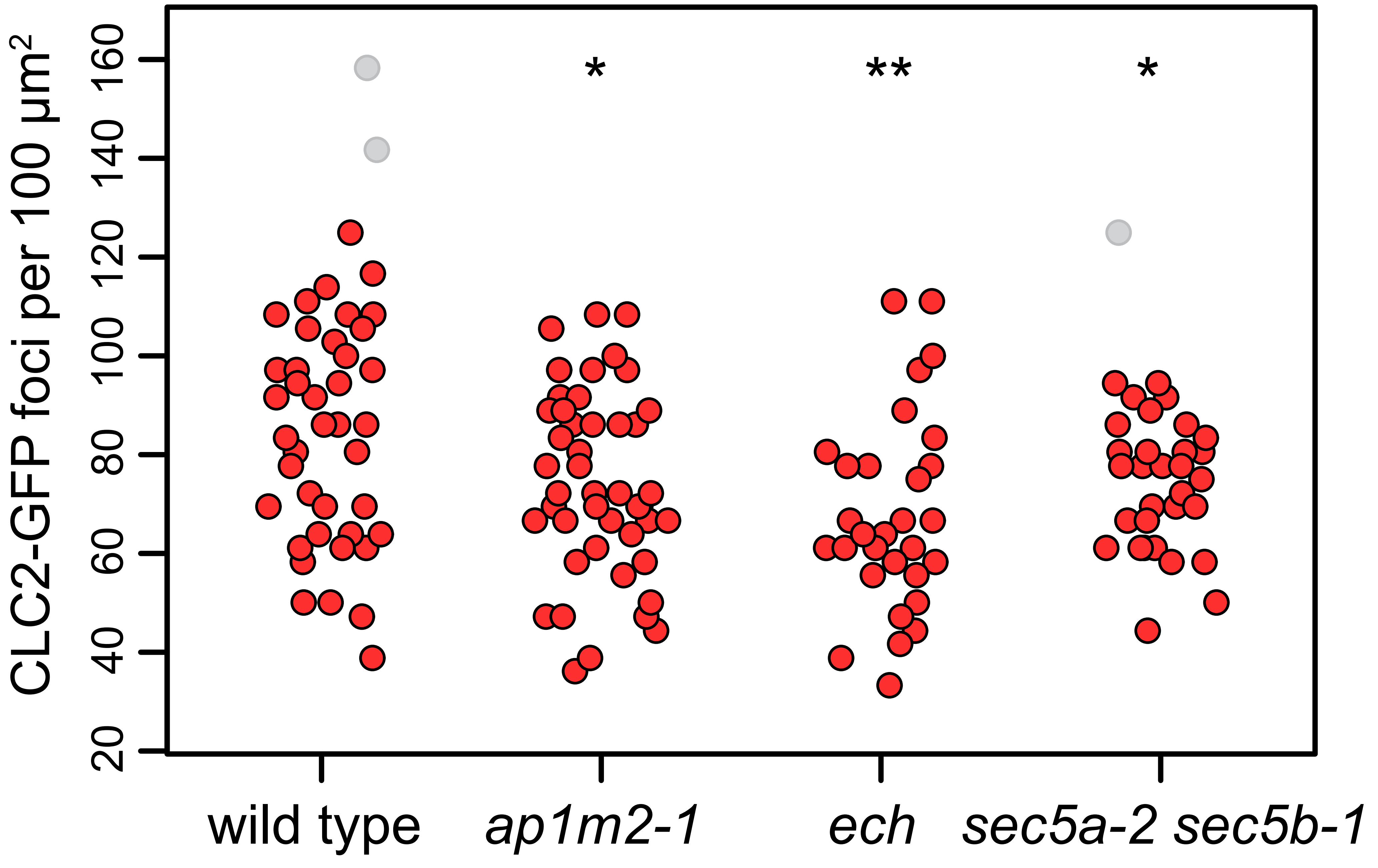


**Suppl. Figure 4. Statistical evaluation of CLC2-GFP foci density after removal of outliers**

Related to Figure 7.

Graph visualising foci density of CLC2-GFP in seedling roots of ap1m2-1, ech, and sec5a-2 sec5b-1 mutants. Each data point represents a measurement from one cell. Data points in grey are outliers of high values removed from this analysis. Mutant values were compared with the wild type control using t tests after removal of outliers, * P≤0.05, ** P≤0.01.

Suppl. Table 1. Lines generated as part of this study

| Line name | Notes |
|---|---|
| *secRFP XVE»amiCHCa* | |
| *secRFP XVE»amiCHCb* | |
| *secRFP XVE»amiCHCc* | |
| *CLC2*$_{pro}$*:CLC2-GFP UBQ10*$_{pro}$*:mCh-AUXILIN-LIKE1 XVE»amiCHCa* | |
| *LAT52*$_{pro}$*:TPLATE-GFP RPS5A*$_{pro}$*:AP2A1-TagRFP tplate XVE»amiCHCa* | |
| *DRP1C*$_{pro}$*:DRP1C-GFP XVE»amiCHCa* | |
| *BEN3*$_{pro}$*:BEN3-TagRFP XVE»amiCHCa* | |
| *ARFA1c*$_{pro}$*:ARFA1c-GFP XVE»amiCHCa* | |
| *GNOM*$_{pro}$*:GNOM-LIKE1-GFP XVE»amiCHCa* | |
| *secRFP HS*$_{pro}$*:PIN1-GFP XVE»amiCHCa* | |
| *secRFP HS*$_{pro}$*:TMK4-GFP XVE»amiCHCa* | |
| *secRFP XVE»AUXILIN-LIKE1* | |
| *HS*$_{pro}$*:PIN1-GFP XVE»AUXILIN-LIKE1* | |
| *HS*$_{pro}$*:TMK4-GFP XVE»AUXILIN-LIKE1* | |
| *ech CLC2*$_{pro}$*:CLC2-GFP* | SAIL_163_E09 |
| *ech LAT52*$_{pro}$*:TPLATE-GFP RPS5A*$_{pro}$*:AP2A1-TagRFP* | SAIL_163_E09 |
| *sec5a-2 sec5b-1 CLC2*$_{pro}$*:CLC2-GFP* | GK-731C01;<br>SALK_001525 |
| *sec5a-2 sec5b-1 LAT52*$_{pro}$*:TPLATE-GFP RPS5A*$_{pro}$*:AP2A1-TagRFP* | GK-731C01;<br>SALK_001525 |
| *ap1m2-1 CLC2*$_{pro}$*:CLC2-GFP* | FLAG_293C11 |
| *ap1m2-1 LAT52*$_{pro}$*:TPLATE-GFP RPS5A*$_{pro}$*:AP2A1-TagRFP* | FLAG_293C11 |

Suppl. Table 2. Primers used in this study

| Primer name | Sequence | Purpose |
|---|---|---|
| ech-F | acagccattgtcgtcctctt | *ech* genotyping |
| ech-R | TCATTGATCTCGTTCCACCA | |
| ap1m2-1-F | tttggtgctggcagTATGAG | *ap1m2-1* genotyping |
| ap1m2-1-R | cagaaaaggaaagtggcaca | |
| sec5a-2-F | AGGAAGATGATGGTGCTTGG | *sec5a-2* genotyping |
| sec5a-2-R | CAAATCTGCTGCAGTCGTGT | |
| sec5b-1-F | ctctttggttgtggcagTCA | *sec5b-1* genotyping |
| sec5b-1-R | TTTGCCAACTCTCTCGCTTT | |
| LB-FLAG | CTACAAATTGCCTTTTCTTATCGAC | |
| LBb1.3 | ATTTTGCCGATTTCGGAAC | |
| LB-SAIL-3 | TAGCATCTGAATTTCATAACCAATCTCGATACAC | |
| LB-GK_o8409 | ATATTGACCATCATACTCATTGC | |
| CHCa-I | gaTCTCGCAGTACTTACCCACAAtctctctttttgtattcc | amiCHCa cloning |
| CHCa-II | gaTTGTGGGTAAGTACTGCGAGAtcaaagagaatcaatga | |
| CHCa-III | gaTTATGGGTAAGTAGTGCGAGTtcacaggtcgtgatatg | |
| CHCa-IV | gaACTCGCACTACTTACCCATAAtctacatatatattcct | |
| CHCb-I | gaTGCAAAATTTACTGCTCCCTGtctctctttttgtattcc | amiCHCb cloning |
| CHCb-II | gaCAGGGAGCAGTAAATTTTGCAtcaaagagaatcaatga | |
| CHCb-III | gaCAAGGAGCAGTAATTTTTGCTtcacaggtcgtgatatg | |
| CHCb-IV | gaAGCAAAAATTACTGCTCCTTGtctacatatatattcct | |
| CHCc-I | gaTTATTTGACCAGTCTTGGCAGtctctctttttgtattcc | amiCHCc cloning |
| CHCc-II | gaCTGCCAAGACTGGTCAAATAAtcaaagagaatcaatga | |
| CHCc-III | gaCTACCAAGACTGGACAAATATtcacaggtcgtgatatg | |
| CHCc-IV | gaATATTTGTCCAGTCTTGGTAGtctacatatatattcct | |
| attB1-amiR-F | GGGGACAAGTTTGTACAAAAAAGCAGGCTCCCCAAACACACGCTCGGA | amiCHC cloning |
| attB2-amiR-R | GGGGACCACTTTGTACAAGAAAGCTGGGTCCCCATGGCGATGCCTTAAA | |
| attB1-TMK4-F | GGGGACAAGTTTGTACAAAAAAGCAGGCTCAATGGAGGCTCCTAC | TMK4 cloning |

| | | |
|---|---|---|
| attB2-TMK4-Rns | GGGGACCACTTTGTACAAGAAAGCTGGGTCCC GACCATCAGCTGA | |
| attB1-PIN1-F | ggggacaagtttgtacaaaaaagcaggctcga tgattacggcggcggacttctaccac | PIN1-GFP cloning |
| attB2-PIN1-R | ggggaccactttgtacaagaaagctgggtctc atagacccaagagaatgtagtagag | |
| TUB2-qF | AAACTCACTACCCCCAGCTTTG | qPCR |
| TUB2-qR | CACCAGACATAGTAGCAGAAATCAAGT | qPCR |
| PP2AA3-qF | TAACGTGGCCAAAATGATGC | qPCR |
| PP2AA3-qR | GTTCTCCACAACCGCTTGGT | qPCR |
| CHC-qF1 | GTTCAGGCTTGCAAGGAGTA | qPCR |
| CHC-qR1 | TCAGGATCCTCACTCATACTCAA | qPCR |
| CHC-qF2 | CCGTATCTCCTCCAGTTTATCC | qPCR |
| CHC-qR2 | ACCTCCCATTCCTGGCATC | qPCR |

Suppl. Table 3. Constructs generated as part of this study

| Plasmid name | Notes |
|---|---|
| PIN1-GFP-2/pDONR221 | with STOP codon |
| TMK4/pDONR221 | without STOP codon |
| amiRNA-CHCa/pDONR221 | |
| amiRNA-CHCb/pDONR221 | |
| amiRNA-CHCc/pDONR221 | |
| proHS/pDONRP4P1r | |
| UBQ10-XVE»amiRNA-CHCa/pH7m24GW,3 | |
| UBQ10-XVE»amiRNA-CHCa/pK7m24GW,3 | |
| UBQ10-XVE»amiRNA-CHCa/pB7m24GW,3 | |
| UBQ10-XVE»amiRNA-CHCb/pB7m24GW,3 | |
| UBQ10-XVE»amiRNA-CHCc/pB7m24GW,3 | |
| HS:PIN1-GFP-2/pK7m24GW,3 | |
| HS:TMK4-GFP/pK7m34GW | |
